# Representation of semantic information in ventral areas during encoding is associated with improved visual short-term memory

**DOI:** 10.1101/2019.12.13.875542

**Authors:** Bobby Stojanoski, Stephen M. Emrich, Rhodri Cusack

## Abstract

We rely upon visual short-term memory (VSTM) for continued access to perceptual information that is no longer available. Despite the complexity of our visual environments, the majority of research on VSTM has focused on memory for lower-level perceptual features. Using more naturalistic stimuli, it has been found that recognizable objects are remembered better than unrecognizable objects. What remains unclear, however, is how semantic information changes brain representations in order to facilitate this improvement in VSTM for real-world objects. To address this question, we used a continuous report paradigm to assess VSTM (precision and guessing rate) while participants underwent functional magnetic resonance imaging (fMRI) to measure the underlying neural representation of 96 objects from 4 animate and 4 inanimate categories. To isolate semantic content, we used a novel image generation method that parametrically warps images until they are no longer recognizable while preserving basic visual properties. We found that intact objects were remembered with greater precision and a lower guessing rate than unrecognizable objects (this also emerged when objects were grouped by category and animacy). Representational similarity analysis of the ventral visual stream found evidence of category and animacy information in anterior visual areas during encoding only, but not during maintenance. These results suggest that the effect of semantic information during encoding in ventral visual areas boosts visual short-term memory for real-world objects.

---

Our visual environment, at any given moment, is overwhelmingly complex. To cope with the abundance of available information, we rely on selective attention to focus upon one thing at a time, and then visual short-term memory (VSTM) to hold and manipulate relevant information. VSTM is therefore a critical for many everyday tasks. However, it is highly limited in capacity. Only a small number of high-fidelity simple features (e.g., color, orientation) can be maintained in VSTM (Luck & Vogel, 2013); and the number of objects that can be maintained in VSTM is further reduced when trying to maintain more complex objects (Alvarez & Cavanagh, 2004).

The origin of this limited capacity has been debated. One theoretical position is that the key limitation is on the amount of information that can be maintained. During the maintenance of VSTM, there is a sustained, load-dependent activity, particularly in parietal regions (Linden et al., 2003; Todd & Marois, 2004; Xu & Chun, 2006) that asymptotes with increasing memory load. It has been proposed that this activity reflects the amount of information actively maintained in VSTM; thus, it could be that the limit on information that can be stored in VSTM is due to these limited storage resources (McNab & Klingberg, 2008; Todd & Marois, 2005).

Another theoretical position is that the primary bottleneck is the encoding of information into VSTM. According to this view, the amount of information about a stimulus that can be maintained is limited by the amount of information encoded by perceptual regions (Emrich, Riggall, Larocque, & Postle, 2013). It is supported by evidence that encoding strategy has a strong effect on individual differences in VSTM (Linke et al, 2011; Cusack et al, 2009), and on the amount of stimulus-specific information recoverable from sensory regions, often without measurable delay-period activity (Harrison & Tong, 2009; Riggall & Postle, 2012; Serences, Ester, Vogel, & Awh, 2009). Increases in memory load can affect the signal-to-noise ratio in populations of feature-specific neurons (Bays, 2014), resulting in decreased decoding from patterns in sensory cortex (Emrich et al., 2013).

The amount of information that can be stored is also affected by factors other than the number of stimuli to be encoded. For example, the amount of knowledge or familiarity an individual has with a stimulus affects the precision and/or the capacity of stored representations. For example, own-race faces are recalled with greater precision than other-race faces (Zhou, Mondloch, & Emrich, 2018). Similarly, Xie and Zhang (2017) demonstrated that familiarity with Pokemon sped up the rate with which objects were encoded into VSTM, as measured by event-related potentials (ERPs; Xie & Zhang, 2018). Memory for colors is affected by the extent to which they belong to the category labels (Bae, Olkkonen, Allred, & Flombaum, 2015; Hardman, Vergauwe, & Ricker, 2017).

It remains unclear, however, what neural mechanisms support the increased ability to store familiar objects in VSTM. Brady et al (Brady, Störmer, & Alvarez, 2016) used the contralateral delay activity (CDA), an ERP component associated with memory storage, to examine storage for familiar objects compared to simple features, and found a greater CDA amplitude throughout the delay period for familiar stimuli. This finding suggests that regions associated with delay-period activity may be able to recruit additional resources for familiar stimuli (perhaps with the recruitment of additional long-term memory regions). It is unclear, however, how familiarity, driven by semantic information, affects processing during encoding, particularly within sensory regions associated with the maintenance and precision of feature-specific information.

One limitation of studies contrasting VSTM for simple features versus familiar objects is that it is not possible to compare VSTM for familiar and unfamiliar stimuli without accounting for object complexity (i.e., the number of perceptual features). That is, although familiar real-world objects are remembered more accurately, they also tend to have greater complexity. Numerous studies have demonstrated that object complexity tends to decrease memory performance, and decreases the amount of storage-related delay-period activity in the superior intraparietal sulcus and lateral occipital complex (LOC; Xu & Chun, 2006, 2009) reflecting increased storage demands, which reach an asymptote at lower memory loads. Recently, Stojanoski & Cusack (2014) developed a method of warping stimuli that, while reducing the available semantics associated with an object, controlled the effect on the physical complexity of the stimulus as processed by early visual areas. Using this warping method, Veldsman and colleagues (Veldsman, Mitchell, & Cusack, 2017a) demonstrated that less-warped versions of objects exhibited more varied activity, rather than changes in the amount of activity, in a number of regions associated with VSTM, suggesting richer neural representations for the better-remembered, intact objects. However, in the study by Veldsman and colleagues, participants were required to compare different levels of warping within individual objects that were not organized into superordinate classes (e.g., categories) across different levels of warping, precluding the measurement of semantic representations. Without manipulating access to semantic content both by grouping images into superordinate classes, while controlling the level of warping across recognizable and unrecognizable objects, it remains unclear what role semantics plays in improving VSTM and how that changes brain activity.

Consequently, the aim of the current experiment was to examine how semantics affected VSTM precision and capacity and to assess neural representations during encoding and maintenance. To do so, we probed VSTM performance by manipulating semantic information in two ways. First, we used a set of objects that were organized at two levels: basic categories (e.g., cars, food) and superordinate (e.g., animate and inanimate). Second, we controlled access to the semantic content of the objects by maintaining the stimulus complexity (warping levels) constant across objects. We hypothesized that access to semantic information will improve visual shortterm memory by increasing both precision and accuracy. This memory advantage for recognizable objects could be driven by recruiting additional anterior regions along the visual hierarchy (DiCarlo, Zoccolan, & Rust, 2012) by improving representations in early visual areas. Storing additional semantic content may also reduce the maintenance load, resulting in changes in the strength of parietal delay-period activity. However, Veldsman, Mitchell and Cusack (2017) found no evidence for changes in the strength of neural activity across the visual hierarchy for remembered recognizable objects. Another possible mechanism is that better memory for recognizable objects is mediated by distinct patterns of neural representations of the semantic information during encoding and maintenance. Moreover, if changes to the neural representation of recognizable objects underlies improved VSTM, we expected these changes to occur primarily in ventral visual areas, in line with an encoding model of VSTM.

## Methods

### Participants

Twenty-two healthy adult volunteers (age 26+/− 4.72 years; 9 male, 13 female) participated in two scanning sessions (at least 6 days apart). A total of forty-two scanning sessions were acquired (two participant sessions were not acquired due to attrition). All participants had normal or corrected-to-normal vision, with no history of neurological problems. All participants provided written consent as required by the local ethics review board, and were compensated $20/hour for scanning.

### Stimuli and Procedure

We used a set of 96 images, taken from the Hemera image database (Hemera Images: http://www.hemera.com/), divided in 8 categories: faces, birds, fruit, mammals, bikes, tools, shoes, and clothes. The object categories could also be divided into two superordinate classes: animate (living) and inanimate (non-living) objects, which has been shown to reflect human semantic representations (Costanzo et al., 2013). To isolate perceptual features from semantic content we created two sets of “warped” images using diffeomorphic transformations, a method developed by Stojanoski and Cusack (2014), which create smooth, continuous, and invertible images that maintain a one-to-one mapping between the source and transformed space (see Stojanoski & Cusack, 2014, for information about the warping algorithm). Therefore, in this context, “low” warped (recognizable) objects were matched in their basic perceptual properties to “high” warped versions but were deemed to be unrecognizable, which was determined based on perceptual and semantic ratings by 415 participants who completed over 15,600 trials on Amazon’s crowdsourcing platform, Mechanical Turk (Fig. 1). Mean warping levels at which all objects per category were no longer recognizable was used to set the warping level threshold for the “high” warp condition in the current neuroimaging experiment. Warping level for images in the “low” warp condition was set to the maximum level that did not disrupt recognizability (see Fig. 1 for sample images). In both the “high” and “low” warp conditions, 16 parametrically varying versions of each item were created confined within the “high” and “low” warp space, respectively. Distance between adjacent images in the set of 16 were mathematically equivalent within and across warping conditions. That is, the distance between any two neighbouring images in the high warp condition was the same as the distance between any two neighbouring images in the low warp condition.

**Fig 1.**
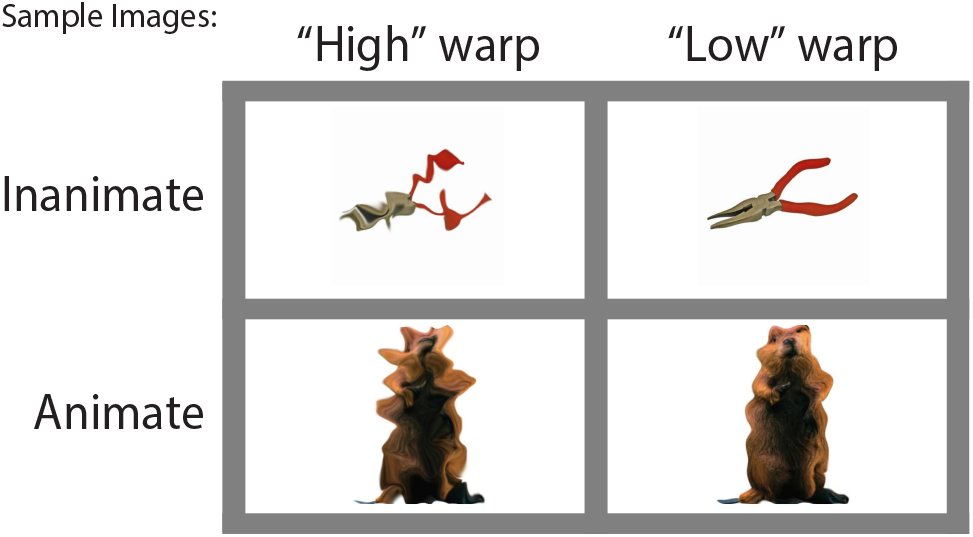
A) A sample stimulus, illustrating the 16 parametrically varying versions of the image, ranging from intact to maximally warped using the diffeomorphic transformations. Images enclosed by the square reflect “low” and “high” warp ratings. These were presented around a circle in the experiment. B) Sample animate and inanimate representing “low” and high” warp conditions.

Participants completed a visual short-term memory task, using a whole report paradigm, while undergoing fMRI. On each trial of a whole report paradigm, participants were required to select the remembered item among a set of distractors presented along a continuous scale. (Fig. 2). The continuous parameter space is modelled to gain an estimate of the precision (the degree of deviation of the reported item from the probed item), and the probability that the item was remembered at all (Wilken and Ma, 2004; Bays and Husain, 2008; Zhang and Luck, 2008). Specifically, their task was to encode, and remember a single image, and identify the remembered item among the set of 16 parametrically varying items positioned along the outline of a response wheel. Target items were randomly selected from the set of 16 versions, with the remaining 15 serving as distractors around the response wheel. At the beginning of each trial a white fixation cross (~20°) was presented in the middle of a gray screen for 1, 6, or 11 seconds, followed by the target item, presented in colour (500 x 500 pixels, 7.9°), which appeared centrally on a gray background, for 3 seconds (Fig 1 b). The offset of the target marked the start of the delay phase, extending 1-11 seconds where participants were instructed to remember the target in as much detail as possible. At the end of the delay period, participants were presented with the response wheel that contained the set of 16 parametrically varying version of the images (image size was reduced to 45 x 45 pixels and were positioned 22.5° apart). With the onset of the response wheel, a black rectangle framed one of the 16 images, at a random location. Participants were instructed to identify the target item by moving the black square (using a MRI compatible button box) until it framed the item that matched with the one in their memory. Participants were given 12 seconds to identify the target; the image inside the frame at the end of the allotted time was taken as their response. Participants completed 96 trials, divided into three runs (32 trials/run) over two scanning sessions, at least 6 days apart. In each session, trials were divided into two types: on half the trials participants saw high warp images and the other half they saw low warp images, with each image presented in both high and low warp conditions, once per session (randomly assigned and counterbalanced across participants). This was designed to avoid perceptual biases a result of initial exposure to low warp images.

**Fig 2.**
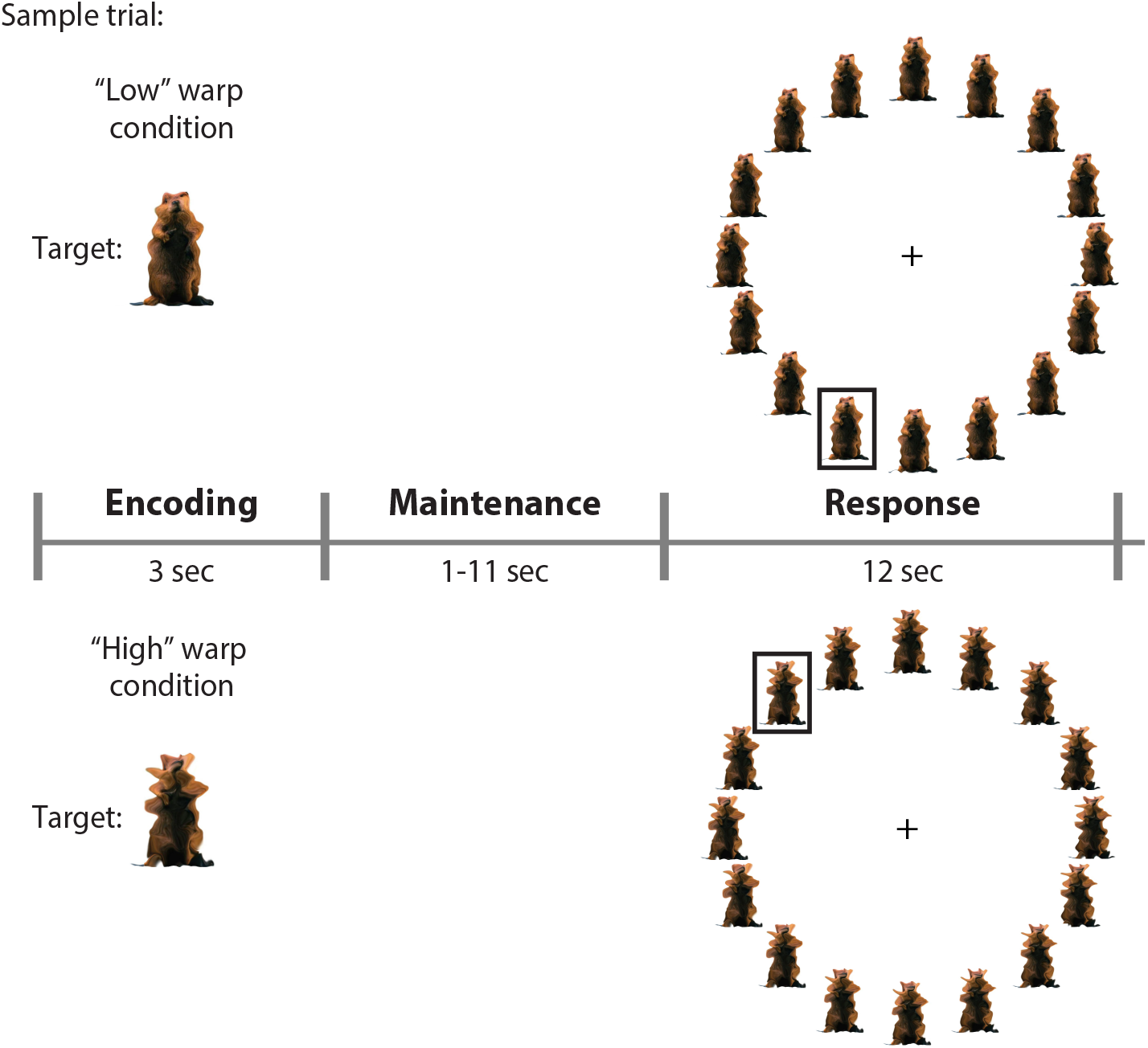
The timeline of two sample trials. At the start of the trial a target item appears in the middle of the screen for 3 seconds, followed by a maintenance period between 1 and 11 seconds. At the start of the response phase a black square surrounds a random distractor, and participants have 12 seconds to move the square until it overlays the item they thing matches the target.

All images were projected (Avotec SV-6011; at 60Hz) onto a screen and were viewed by participants through a mirror mounted on the head coil. MATLAB (MathWorks) and the Psychophysics Toolbox (http://psyctoolbox.org; Brainard, 1997; Pelli & Vision, 1997) were used to deliver stimuli.

### fMRI Acquisition

Participants were scanned with a Siemens Tim Trio 3T MRI scanner. At the start of each scan, a whole brain T1-weighted high-resolution structural image was acquired with an MPRAGE sequence (FOV = 240 x 256, flip angle = 9°, TR = 2300 msec, TE = 2.98 msec, resolution = 1 mm isotropic). Functional images were acquired using a highly accelerated gradient-echo EPI sequence (Center for Magnetic Resonance Research, University of Minnesota) with multiband acceleration factor 3 and GRAPPA iPat acceleration of 2. The following parameters were used: 32 slices were acquired with a matrix size of 70=70 and a voxel size of 3 x 3 x 3 mm (not inclusive of a 10% slice gap), flip angle = 55°, TE = 25 ms, and TR = 850 ms, and a bandwidth of 1587 Hz/Px. Each scanning session was divided into 3 runs, for a total of 129 runs.

## Analysis

### Behavioural Analysis

Behavioural performance on the working memory task was analyzed by first projecting participant’s responses onto a circular distribution ranging from −π < x < π radians with the target at zero, for each trial. Using that information, we calculated the difference between the target position and the response position producing a measure of the degree of error represented as a distribution. The distribution of errors were fit with a probabilistic mixturemodel using the MemToolbox Suchow, Brady, Fougnie, & Alvarez, (2013), to generate maximum-likelihood estimates of precision and guessing rate (Zhang & Luck, 2011). Briefly, the guessing rate is modeled as the height of a uniform distribution, reflecting random responses, whereas precision is estimated as the inverse of the circular normal (Von Mises) distribution on the remaining trials (i.e., those trials in which the target was correctly reported). Due to task-related constraints there were too few trials to fit the model for each participant; instead, we pooled errors across participants to estimate precision and guessing rates across both warping conditions. We also computed the root-mean square (RMS) error, that is the difference between the target and the selected item, for high and low warped objects grouped by category, animacy, and all objects independent of category.

### Imaging Analysis

Functional imaging data was analyzed with SPM8 (Wellcome Institute of Cognitive Neurology; http://www.fil.ion.ucl.ac.uk/spm/software/spm8/), by establishing an analysis pipeline using the automatic analysis system, version 4 (www.github.com/rhodricusack/automaticanalysis). Preprocessing steps in the pipeline followed these six steps: 1) all volumes were converted to Nifti format, 2) motion was corrected by extracting six motion parameters: translation and rotation for three orthogonal axes, 3) brains were normalized, using SPM8 segment-and-normalize procedure where the T1 (anatomical) was segmented into gray and white matter and normalized to a pre-segmented volumetric template in MNI space, 4) extracted normalization parameters were then applied to all function (echo-planar) volumes, 5) data was smoothed using a Gaussian smoothing kernel of 10 mm FWHM (for univariate analyses only; Peigneux et al., 2006), and 6) low frequency noise (e.g., drift) was removed by high-pass filtering the data with a threshold of 1/128 Hz. Four dummy scans at the start of each session were discarded to allow for T1 relaxation.

### Univariate Analyses

We used univariate analyses to identify activation in brain regions, either during the encoding, or maintenance phase of visual short-term memory that varied with level of recognizability (high warp vs. low warp images) in general, between object categories, or animacy. We did this by fitting a general linear model (GLM) to the functional imaging data with separate regressors for high and low warped images for each category during the encoding and maintenance stages of visual short-term memory. Regressors comprised the onsets and durations of each event: during the encoding phase, onsets were defined as the time when the images appeared on the screen, and duration was set to the time the image remained on the screen (3 sec). The onset of the maintenance phase was marked by a white plus sign in the middle of the screen, and duration was the period of time participants were asked to hold the target item in memory (1 – 11 seconds). These time courses were convolved with the canonical hemodynamic response function supplied by SPM. The random jitter ITIs served as a baseline. Contrasts were established to compare encoding and maintenance of high and low warp images versus baseline, and to directly compare high versus low warp images during encoding and maintenance. All results were corrected for multiple comparisons at p < 0.05 FWE.

### Multivariate Analyses: Representational similarity analysis

In addition to examining whether the availability of semantic content (low warp images) resulted in an increase of brain activity in certain brain regions, or recruited different brain regions, we used multivoxel pattern analysis (MVPA) to determine whether representations in visual and parietal regions differed during encoding and maintenance of high and low warped images. We focused our multivariate analyses on four regions of interest (ROIs): bilateral calcarine sulcus, superior parietal cortex, the fusiform area as defined in the AAL atlas (Tzourio-Mazoyer et al, 2002) using the MarsBar ROI package Brett, Anton, Valabregue, & Poline, (2002) as well as the lateral occipital cortex ROI (8 mm sphere around [43, −67, −5] on the right, and [−41, −71, −1] on the left) used by Xu and Chun (2006). We selected these ROIs because they have been shown to be involved in object category processing, and the a priori hypothesis that semantic information would be represented in anterior visual areas during maintenance (i.e., Lateral Occipital Cortex; Todd & Marois, 2004; Xu & Chun, 2006) and encoding (i.e., fusiform gyrus; Connolly et al., 2012; Huth, Nishimoto, Vu, & Gallant, 2012) but not in early visual areas (i.e., bilateral calcarine sulcus) which is mainly linked to encoding and maintenance of simple perceptual features (Christophel, Hebart, & Haynes, 2012). We included a “memory” ROI that was extracted from the univariate analysis during maintenance period for use in the MVPA analysis of the maintenance period. This way, we could assess both global signal changes as well as potential representational differences in regions most sensitive during maintenance. All ROIs remained in normalized space, and all data was gray matter masked for the multivariate analysis. Within these specific ROIs we used MVPA to examine the neural representations of semantic content across our ROIs during the encoding and maintenance phases of visual short-term memory. Specifically, we used representational similarity analysis (RSA), a correlation-based approach that is insensitive to modulations in mean magnitude activations. We fit the data with the same GLM with individual regressors for high and low warp objects for all categories during encoding and maintenance as we used for the univariate analysis. To mitigate the effects of comparisons across different temporal distributions, we confined our comparisons across runs, and only during encoding and maintenance. Beta values for each participant and all events were extracted for each voxel in our ROIs and were Spearman correlated within and across runs. Correlations were normalized to ensure that each run contributed equally. The result of averaging correlations across runs produced a 48 x 48 (12 conditions, 2 warping levels, 2 phases) similarity matrix which was contrasted by warping, animacy, and category matrices for both encoding and maintenance using a GLM (figure for result).

For the warping contrast, images were grouped together based on level of warping. This contrast tested whether the patterns of activity produced by the same warping level (high or low) were more similar to one another than repetitions of the opposite warping level – is the pattern of activity produced by low warp images distinct from the patterns produced by high warp images. We grouped images according to animacy (defined by Kriegeskorte et al., 2008) to run the animacy contrast to test whether activity patterns within animate objects differed from activity patterns produced by inanimate objects for both warping levels. Finally, we ran a category contrast; images were collapsed into semantic categories and tested whether patterns of activity were more similar within a category than activity across categories at both high and low warping levels. Differences emerging in the latter two contrasts would suggest specific ROIs represent either the lower-level properties of the image or their semantic properties. All results were corrected for multiple comparisons using Bonferroni correction.

## Results

### Behavioural Results

As a first method to assess performance, we computed the root-mean square (RMS) error between the target and selected item across each participant’s responses. A two sample t-test showed that for recognizable objects the errors were significantly less distant from the target compared to unrecognizable objects (t(41) = 2.85; p = 0.007; Cohen’s d = 0.44; BF10 = 5.61). To test how memory was better, we fitted the response distributions using a probabilistic mixture model, which gave separate estimates of guessing (i.e., item completely forgotten) and precision (i.e., less accurate memory). The results are shown in Fig. 3. Participants guessed more in the high warp condition, using the Kolmogorov-Smirnov test non-parametric test to compare prior probability distributions (KS = 0.96, p<0.0001) and showed lower precision (KS = 0.49, p < 0.0001).

**Fig 3.**
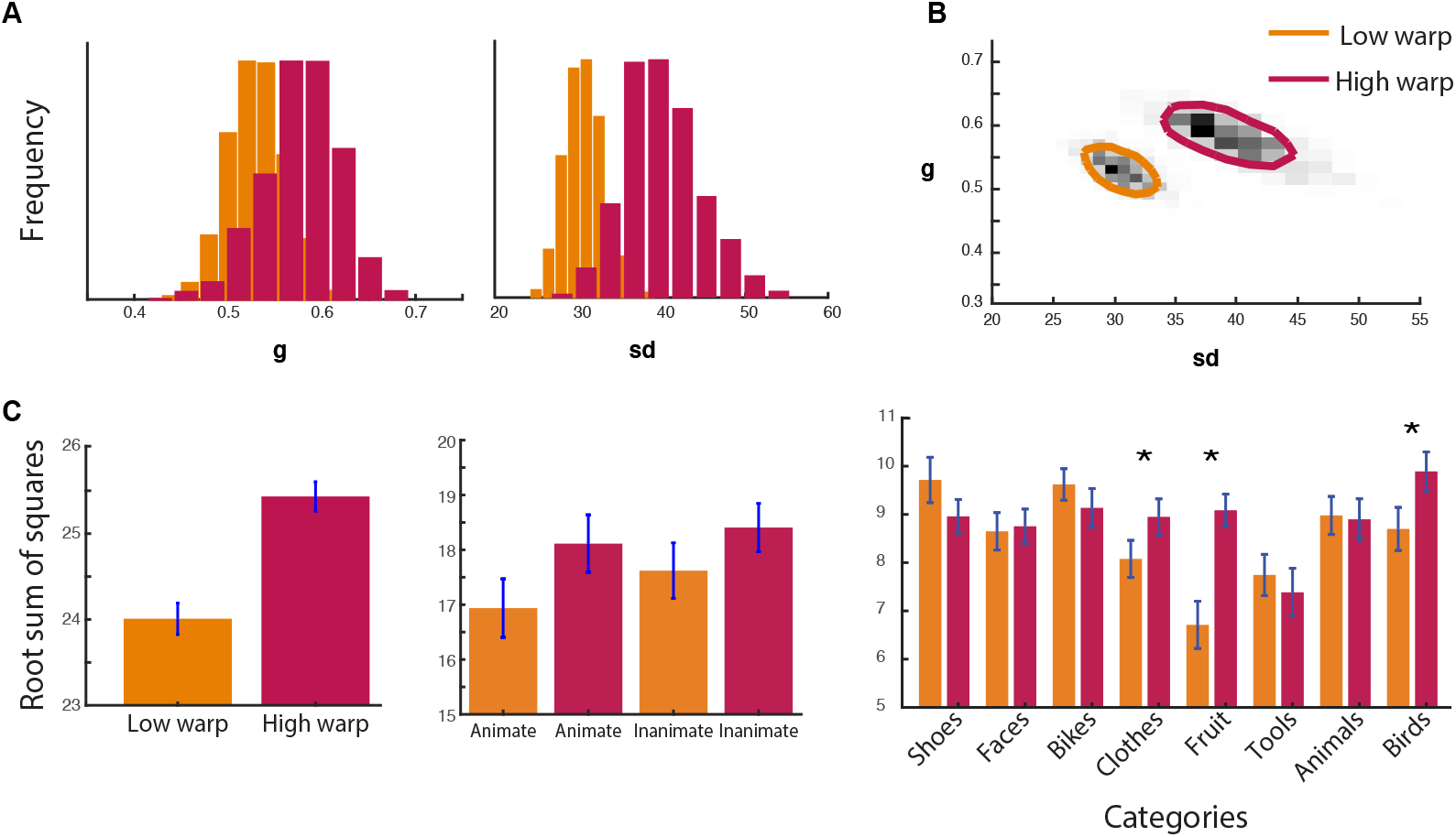
A) The marginal posterior probabilities of the standard mixture model (Suchow et la., 2013) for guessing rate (*g,* left) and the variance of participants’ response around the target item (*sd,* right). Recognizable low warp items have a lower guessing rate and are represented more accurately. B) The joint distribution of guessing rate (*g*) and variance (*sd*). (C) The root-mean square error (RMS) of the target and selected item across each participant’s responses for high and low warped objects (left); animate and inanimate objects (middle); and each object category (right).

We also compared memory performance (distance between target and response) for objects grouped at the level of animacy and category. At the level of animacy, we ran a two-way ANOVA (Recognizability [low warp, high warp] x Animacy [Animate, Inanimate]) and found only a main effect of recognizability (F(1,41) = 9.66; p = 0.003; n2 = 0.19; BF10 = 6.46), and no main effect of animacy or an interaction between recognizability and animacy. This result suggests that memory performance was better for recognizable objects independently of whether those objects were animate or inanimate. At the category level, we also ran a two-way ANOVA (Recognizability [low warp, high warp] x Category [faces, birds, fruit, mammals, bikes, tools, shoes, and clothes]), and found a main effect of Recognizability (F(1,41) = 8.49; p = 0.0006; n2 = 0.17; BF10 = 1.37), a main effect of category (F(1,41) = 5.76; p = 2.11e-5; n2 = 0.12; BF10 = 8503), and a Recognizability x Category interaction (F(1,41) = 5.18; p = 5.88e-5; n2 = 0.11; BF10 = 772.38). These results indicate that overall participants better remembered recognizable objects relative to unrecognizable objects across all categories, but certain recognizable categories were more memorable than others (Fig. 3). Together, we found that semantic information helps in remembering objects in visual short-term memory, likely by increasing both the number of visual features stored in visual short-term memory and the precision of those memories.

### fMRI results

#### Univariate: Whole brain results

Figure 4 shows the pattern of activity during the encoding and maintenance of the various recognizable and unrecognizable objects in our image set. As seen in the top panel, activity during encoding of recognizable objects was associated with fronto-parietal network (Linke, Vicente-Grabovetsky, Mitchell, & Cusack, 2011), occipital and ventral stream regions, including the fusiform area, replicating previous findings examining encoding of real-world objects (Veldsman, Mitchell, & Cusack, 2017). A similar network of regions were activated during the encoding phase of unrecognizable objects. During the maintenance period (in the absence of visual stimulation), however, significant activity was largely limited to the early visual cortex for both recognizable and unrecognizable objects. Reflecting their similarity, we found no difference in the neural activity evoked by recognizable vs unrecognizable objects during either the encoding or the maintenance period. This suggests that processing recognizable objects relies on the same set of brain regions as processing unrecognizable objects, and provides no support for the hypothesis that recognizable objects recruit additional brain areas.

**Fig 4.**
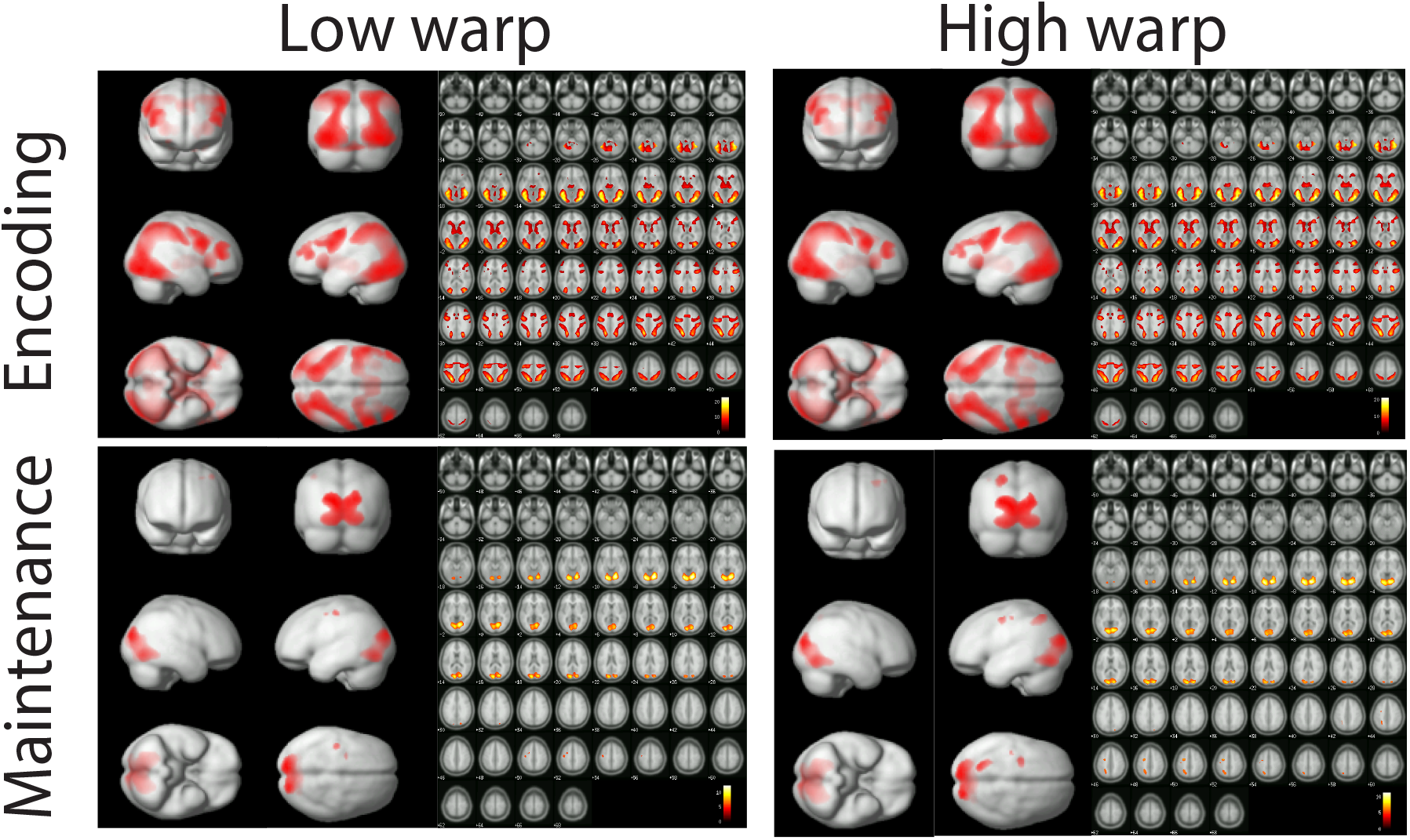
Activity during encoding (top) and maintenance (bottom) of low-warp objects (left) and high warp-objects (right). No effect of the degree of warping on brain activity was found. Colour bars represent t-values. All contrasts are relative to implicit baseline. FDR<0.05.

#### Representational Similarity Analysis: ROI results

Perhaps the memory advantage for recognizable objects was due to differences in the pattern of neural activity, rather than in the overall strength or distribution of neural activity. In other words, is semantic information associated with the recognizable objects represented in distinct patterns of neural activity? To test this hypothesis, we ran a representational similarity analysis (RSA) to compare the similarity of the patterns of neural activity (across repetitions) organized across three levels of semantic information: recognizability (amount of warping), category and animacy, within four ROIs of interest during both encoding and maintenance.

#### Encoding

During encoding, we first compared whether patterns of neural activity are best fit by a model representing recognizability irrespective of category (i.e., recognizable vs. unrecognizable images). The results of the RSA revealed that the pattern of neural activity in response to recognizable and unrecognizable objects did not differ in any of the of the ROIs (t < 1.09; p > 0.05), nor did we find representations differ between ROIs [F(3,123) = 0.44; p < 0.72; n2 = 0.11; BF10 = 0.05]. That is, the brain did not produce a distinct pattern of activity that differentiated recognizable from unrecognizable objects, across the various categories in the regions we selected. This reflects the fact that the warping method we used (Stojanoski & Cusack, 2014) successfully preserved the perceptual properties for both recognizable and unrecognizable objects.

However, we did find evidence for representation of semantic content in the form of animacy and category membership. We examined whether patterns of neural activity in each of the ROIs matched a model that represented animacy (i.e., recognizable animate vs. inanimate objects), the results of the RSA revealed that the fusiform gyrus (t = 2.85; p = 0.007) and the LOC (t = 3.17; p = 0.003), but not the other ROIs, produced distinct neural representations for animate and inanimate objects. We also found that the representations of animacy for recognizable objects was significantly stronger than that for unrecognizable objects within both the fusiform gyrus (t = 2.07; p = 0.045; Cohen’s d = 0.32; BF10 = 1.15) and the LOC (t = 2.42; p = 0.02; Cohen’s d = 0.37; BF10 = 2.21). We found a similar pattern of results for category information. That is, the pattern of neural activity matched a model representing category membership in the LOC (t = 2.92; p = 0.006), but the model fit was not significant in the other ROIs (after Bonferroni correction). This effect was also significantly stronger than patterns of neural activity representing category information for unrecognizable objects in LOC (t = 2.35; p = 0.024; Cohen’s d = 0.37; BF10 = 1.94). Together these results suggest that semantic information is extracted primarily in the fusiform gyrus and LOC, while this information cannot be decoded in earlier visual areas or in the parietal cortex (Fig. 5).

**Fig 5.**
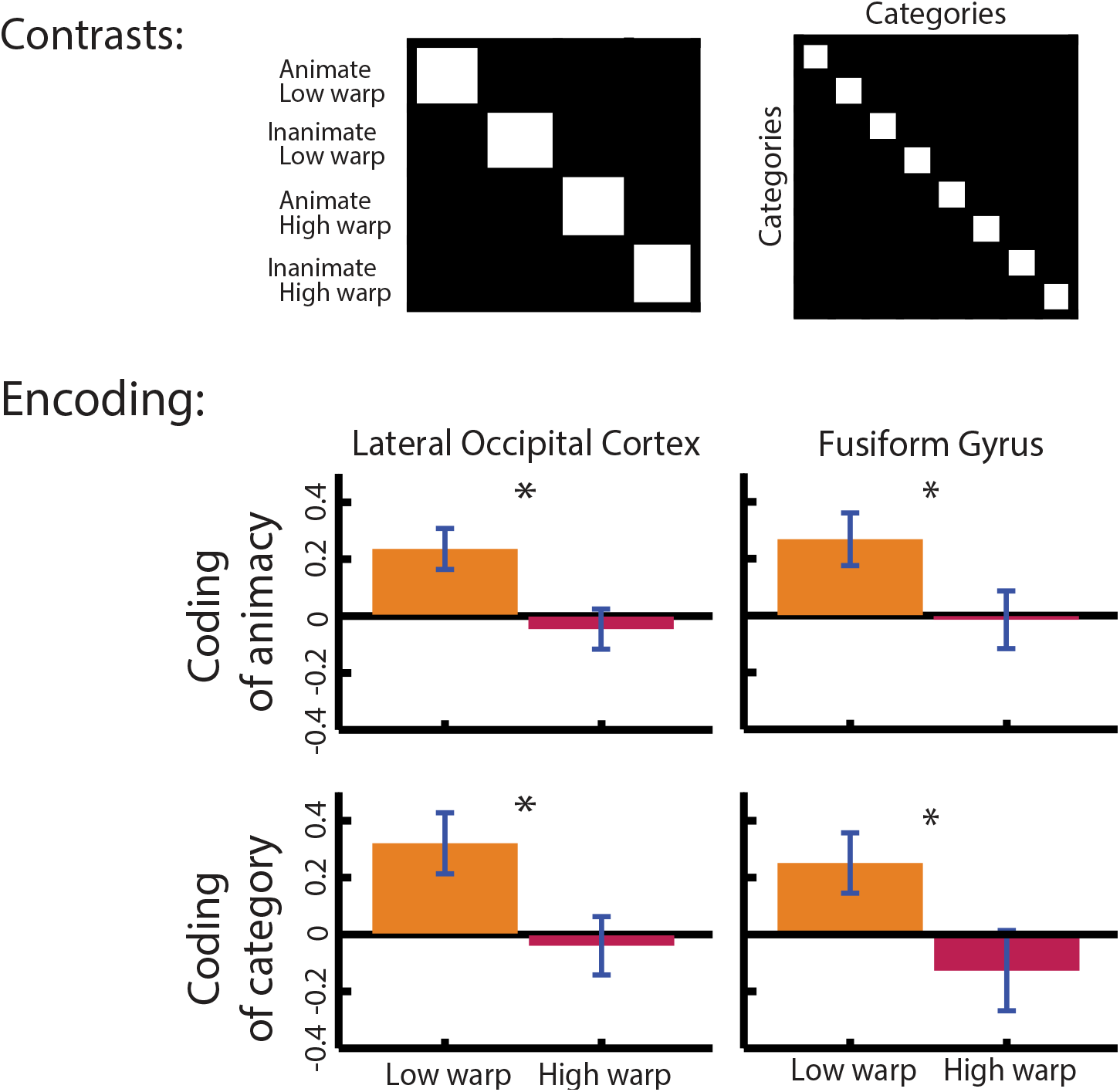
Top panel: The models used to conduct the representational similarity analysis. Lower panel: Beta values produced by the general linear model used to summarize the representational similarity analysis. Results depict differences between low and high warp in the Lateral Occipital Cortex, and the Fusiform Gyrus

#### Maintenance

To assess whether semantic information about the objects is also present during maintenance we conducted the same RSA analysis described above. Much like during encoding, we found no evidence that patterns of neural activity differed between recognizable from unrecognizable objects within any of the ROIs (t_(Bonferroni corrected)_< 2.27; p > 0.12). We also examined whether neural representations for recognizable and unrecognizable objects differed between ROIs, but we found no significant differences (F(4,164) = 1.06; p = 0.37; n2 <0.03; BF10 = 0.07). However, unlike during encoding, we found no evidence that semantic information was encoded during maintenance. That is, we did not find distinct patterns of activity in any of the ROIs that represented animacy (t <1.99; p >0.053), or category membership (t <1.97; p >0.056) in any of the ROIs. A three-way repeated measures ANOVA (Recognizability [low warp, high warp] x Identity [Category, Animacy] x ROI [Calc, LOC, FF, Par]) did not reveal any significant main effects or interactions (F(1,41) < 1.73; p > 0.2; n2 < 0.41; BF10 < 1.12), aside from a significant Identity x ROI interaction (F(3.2,133.2) = 3.35; p = 0.014; n2 < 0.08; BF10 = 6.77), reflecting the fact that animacy for both recognizable and unrecognizable objects was encoded more strongly in parietal cortex (and no other ROI) relative to category membership. In sum, semantic information was not represented during maintenance despite this information being encoded during the perception stage of the visual short-term memory task.

## Discussion

The aim of the current study was to examine the role of semantic information about real-world objects on neural measures of visual short-term memory. We used a novel warping method (Stojanoski & Cusack, 2014) that distorts intact objects in a manner that preserves perceptual features of objects while making them unrecognizable. In this way, we could tease out the influence on semantic content on visual short-term memory performance, as well as the underlying neural mechanisms, without affecting the low-level properties associated with those stimuli.

We found that low-warped images, with intact semantic content, were remembered better than high-warped objects that could not be recognized. By calculating target selection using a continuous report paradigm and a mixture model we found the memory benefit for recognizable objects was reflected in both more precise memory and a lower guessing rate. Moreover, we also found this memory benefit for objects grouped by both animacy and category: both recognizable animate and inanimate objects were remembered better than unrecognizable animate and inanimate objects. Similarly, recognizable objects clustered into basic-level categories were remembered with more precision than clustering of the same categories of unrecognizable objects. These findings suggest that various forms of semantic information are incorporated in visual short-term memory representations that help boost memory performance.

What are the neural mechanisms that support semantically driven improvement to visual shortterm memory? To address this question, we examined changes in brain activity during the encoding and maintenance periods of visual short-term memory. The results of our whole-brain univariate analyses revealed that the encoding period was associated with activity in frontoparietal network (Linke, Vicente-Grabovetsky, Mitchell, & Cusack, 2011; Stokes, 2015), occipital and ventral stream regions, such as the fusiform gyrus, while activity was restricted primarily to early visual cortex during maintenance. Importantly, this pattern of neural activity during encoding and maintenance was the same for recognizable and unrecognizable images; we found no differences in the strength of brain activity and no recruitment of distinct brain regions.

If no additional activity or regions were observed for recognizable compared to scrambled objects, what can account for the behavioural improvements? An RSA analysis revealed that during encoding, but not during maintenance, semantic content representing category and animacy information could be decoded from patterns of activity in the fusiform gyrus and LOC. However, the neural representations associated with category and animacy was not present during the maintenance phase. This finding suggests that it is the extraction of semantic information during encoding by higher ventral stream visual areas that allows these objects to be encoded with greater detail. Importantly, this effect was not observed in early visual areas. Thus, semantic information was restricted to those regions that process information about object categories and identities (Barense, Gaffan, & Graham, 2007; Barense, Henson, Lee, & Graham, 2010; Tyler et al., 2013) and cannot be attributed to differences in low-level featural information.

Although we did not find evidence that semantic information was represented in sensory regions during the delay period, past studies have found evidence for this effect. For example, Lewis-Peacock and colleagues (Lewis-Peacock, Drysdale, & Postle, 2015) used multi-voxel pattern analysis to decode the semantic dimensions of visual stimuli. However, this activity was primarily evident when the semantic (as opposed to visual or verbal) content of the image was task-relevant. Thus, it’s possible that because the task did not require participants to use the semantic content in the task, this activity was absent from the delay period, consistent with findings that VSTM representations can change across tasks (Vicente-Grabovetsky, Carlin, & Cusack, 2014). Nevertheless, the finding that performance was better for the low-warped images suggests that the obligatory coding of semantic information during encoding confers a memory advantage, even if semantic information is irrelevant to completing the task.

It is also possible that information about semantics continues to exist in ventral visual areas during the delay period, but in an “activity silent” state. That is, recent studies, have demonstrated that stimulus and category specific representations can be recovered from sensory areas, even when it is not immediately apparent in the delay activity (Rose et al., 2016; Stokes, 2015). The recovery of these representations in the absence of ongoing activity suggests that this information might be stored in a latent state, perhaps through synaptic weights (but see Schneegans & Bays, 2017). Thus, it is possible that semantic information continued to be represented but was not recoverable from the ongoing activity alone, perhaps because that information was no longer in the focus of attention during the delay period.

While the contribution of semantics has been studied extensively in other domains, such as long-term memory (Hollingworth & Henderson, 1998) and attention (de Groot, Huettig, & Olivers, 2016), understanding how semantics influences visual short-term memory is still at the incipient stages. Our results indicate that semantic information about category and animacy membership plays an important role in visual short-term memory for real-world objects. This is in-line with a growing body of evidence supporting the notion that semantics can influence various aspects of working/short-term memory. For instance, O’Donnell, Clement, & Brockmole, (2018) argue that semantic information increases the capacity of visual working memory, by showing improved memory for image arrays containing semantically related interacting objects (i.e., a key and a lock). Moreover, Veldsman, Mitchell, and Cusack, (2017) showed that the precision of visual short-term memory improves when comparing memory performance for recognizable versus unrecognizable objects. Extending their findings, we show that it is not only the semantics associated with individual objects, but also semantic information about animacy and category inclusion that increases visual short-term memory performance.

This introduces a potential paradox: real-world objects are more “complex” than simple features, such as a colour patch, and complexity is typically associated with a decrease in working memory capacity (Xu & Chun, 2006, 2009), yet, we found memory performance was better for recognizable objects. The warping method used here allowed us to hold visual complexity constant, while preserving semantic information only for the low-warp images. Thus, while real-world objects may contain more visual complexity than simple features, access to semantic information to similarly complex objects boosts memory performance. One way semantic information may help to reduce memory load is by allowing for objects to be encoded at an abstracted level (Christophel, Klink, Spitzer, Roelfsema, & Haynes, 2017), which provide a type of schema that make changes between features more apparent. For example, both neuroimaging studies and behavioural modelling have demonstrated a memory advantage for colors that are easily put into color categories compared to those which require fine-scaled discriminations (Bae, Olkkonen, Allred, & Flombaum, 2015; Hardman, Vergauwe, & Ricker, 2017; Lara & Wallis, 2014). This is consistent with past studies examining memory for up-right versus inverted faces (Lorenc, Pratte, Angeloni, & Tong, 2014). Similarly, Zhou et al (2018) have shown that with short exposures, VSTM for own-race faces was better than for other-races faces, suggesting that stimulus familiarity sped the rate of encoding for familiar own-race faces. This idea is consistent with a couple of mechanisms underlying a semantically driven boost in memory that have recently been proposed. For instance, O’Donnell, Clement, & Brockmole (2018) and Curby, Glazek, & Gauthier (2009) have suggested that access to the semantic properties of objects limits processing resources, allowing them to be more efficiently represented, and thereby increasing working memory capacity. Whereas, Veldsman and colleagues (2017) showed that a richer and wider range of neural representations supports improved visual short-term memory for real-world objects.

What is common between these proposed mechanisms is that benefits to visual short-term memory arise at encoding and not during maintenance, which is consistent with an encoding account of visual short-term memory. Importantly, our results are also consistent with an encoding mechanism, as no differences were observed during the maintenance period in either the univariate analysis or the RSA analysis. In other words, although more information was encoded about intact objects, maintaining that information did not require additional activity or the recruitment of additional brain areas. This is in contrast to some past studies that have demonstrated greater maintenance-related activity for real-word objects compared to simple features (Brady et al., 2016; Galvez-Pol, Calvo-Merino, Capilla, & Forster, 2018; Wong, Peterson, & Thompson, 2008). However, given that these past studies did not control for the complexity of the stimuli, it is possible that it is the greater object complexity, rather than the semantic information per se, that was driving this effect. Consequently, our finding underscores the importance of having appropriately matched stimuli in order to properly dissociate the effects of complexity from the contributions of semantic information to neural measures of VSTM.

## Acknowledgments

BS and RC were supported by NSERC Discovery grant (150688084). SME was supported by NSERC Discovery grant (2019-435945).

